# Brain reorganization in anticipation of predictable words

**DOI:** 10.1101/101113

**Authors:** Jeremy I Skipper, Jason D Zevin

**Affiliations:** Experimental Psychology, University College London, London, UK; Departments of Psychology and Linguistics, University of Southern California, Los Angeles, CA, USA; Haskins Laboratories, New Haven, CT, USA

## Abstract

How is speech understood despite the lack of a deterministic relationship between the sounds reaching auditory cortex and what we perceive? One possibility is that unheard words that are unconsciously activated in association with listening context are used to constrain interpretation. We hypothesized that a mechanism for doing so involves reusing the ability of the brain to predict the sensory effects of speaking associated words. Predictions are then compared to signals arriving in auditory cortex, resulting in reduced processing demands when accurate. Indeed, we show that sensorimotor brain regions are more active prior to words predictable from listening context. This activity resembles lexical and speech production related processes and, specifically, subsequent but still unpresented words. When those words occur, auditory cortex activity is reduced, through feedback connectivity. In less predictive contexts, activity patterns and connectivity for the same words are markedly different. Results suggest that the brain reorganizes to actively use knowledge about context to construct the speech we hear, enabling rapid and accurate comprehension despite acoustic variability.

---

A long history of lexical priming studies in psychology demonstrates that hearing words activates associated words^1^. For example, in a lexical decision experiment, the prime ‘pond’ results in faster reaction times to the subsequent presentation of ‘frogs’, compared to unrelated words. Whether explained in terms of spreading activation among semantically related words^2^, and/or generative prediction^3,4^, primes may serve as part of a solution to the problem of how humans so easily perceive speech in the face of acoustic variability. Despite more than 50 years of searching, speech scientists have found *no* consistent acoustic information that can account for perceptual constancy of speech sounds^5,6^. Primed words might help mitigate this problem by serving as *hypotheses* to test the identity of upcoming speech sounds, thereby constraining interpretation of those indeterminate or ambiguous patterns as specific categories^6–9^. For example, the sentence context ‘The pond was full of croaking…’ primes ‘frogs’. This can serve as an hypothesis to test whether there is enough evidence to interpret the following acoustic pattern as an /f/ despite the uniqueness of that particular ‘f’.

We propose that the neural implementation of this ‘hypothesis-and-test’ mechanism involves the ‘neural reuse’^10^ of processing steps associated with speech production^6,11^’^12^. These steps, ‘selection’, ‘sequencing’, ‘prediction’ and ‘comparison’, are implemented in a sensorimotor speech production network. When someone wants to speak, a word must first be selected from among competing alternatives say (e.g., ‘frogs’ over ‘toads’). The selection is then sequenced into a series of articulatory movements associated with smaller speech units (e.g., /f/, /r/, etc.). Through feedback or ‘efference copy’, the brain predicts the sensory goals of these vocalizations^13^, increasing sensitivity to acoustic patterns related to predicted outcomes in auditory cortex (AC)^14,15^. This allows for vocal learning and continuous adjustment of vocalizations in real time^16,17^. For adjustments to occur, the predicted sensory goal must be compared to the actual information arriving in AC. If an ‘error signal’ is generated, forward and backward propagation continue in the speech production network until it is suppressed (i.e., until adjustment is achieved).

Building on theories of speech perception that posit a central role for motor representations, we hypothesize that a parallel set of operations is reused to achieve perceptual constancy during speech perception (Figure 1). As priming studies demonstrate, listening context results in the activation of related though unpresented words. To serve as an hypothesis, the word with the most activation is implicitly selected and sequenced. This occurs as in speech production but without overt vocalization. The subsequent production steps, prediction and comparison, make sense of why these steps facilitate perceptual constancy. Specifically, sequenced speech units activate the sensory goals corresponding to predicted words. Those goals are ‘tested’ by comparing them to signals arriving in AC. When the hypothesis is accurate, processing demands are reduced because no ‘error signal’ is generated and feedforward and feedback propagation need not occur or continue.

**Figure 1.**
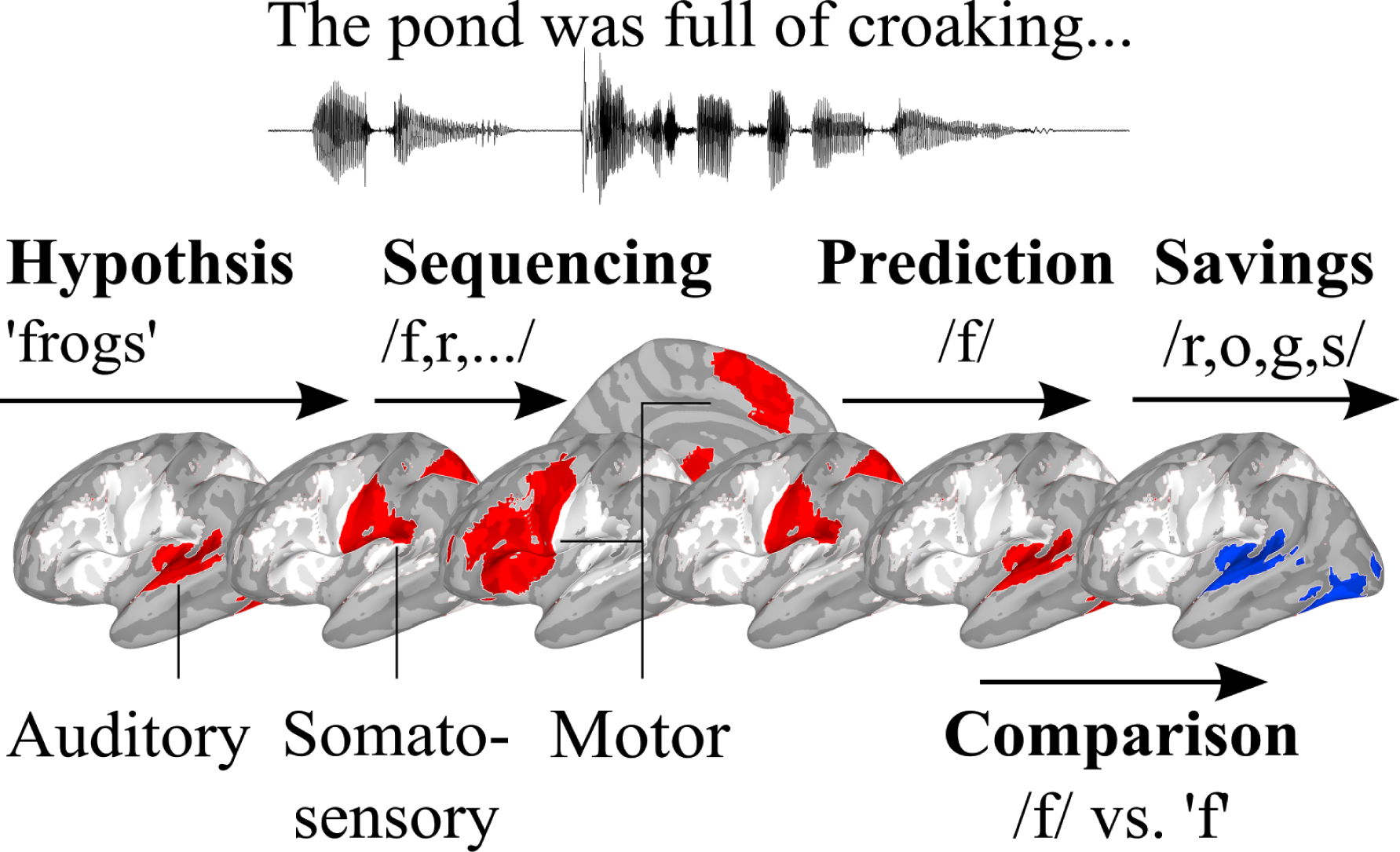
Hypothesis-and-test model of speech perception. Sentence context results in the activation of associated words that serve as hypotheses about subsequent sounds. Brain regions for producing oral/facial movements (red/white/blue) are used to ‘test’ hypotheses. The hypothesis with the most activation (1 red) is selected and sequenced into segments, involving parietal/somatosensory (2 red) and pre- and primary motor cortices (3-4 red). This results in a prediction of the somatosensory/acoustic consequences of producing those segments through feedback (5-6 red). Predictions are compared to arriving auditory information, with processing savings when accurate compared to less constraining contexts (7 blue).

Consistent with this proposal, behavioral evidence suggests that speech production systems are involved in making use of sentence context. For example, hearing “The pond was full of croaking…” influences tongue position, affecting the subsequent forced production of a word (like “toad”) compared to “frogs”^18^. Generally, prediction appears to play a role in the neurobiology of perceiving speech^11^’^19–21^. Furthermore, speech production associated networks seem to play ubiquitous roles in speech perception^22^ and more predictable speech results in a decrease in AC activity compared to less predictable speech^11^. However, inferences about the role of prediction in the neurobiology of speech perception have been based entirely on differences between responses to more versus less predictable words once they have been presented. Thus, the general mechanisms supporting prediction have not been observed. And, though it has long been hypothesized that speech production processes are important for dealing with the indeterminacy of the speech signal^7,23^, and contemporary models of speech perception increasingly incorporate an active hypothesis-and-test like mechanism^6,24–26^, this specific mechanism has also never been observed as it would be engaged before predictable words.

Thus, we tested the described hypothesis-and-test model in two functional magnetic resonance imaging (fMRI) studies, one involving experimental manipulation of sentences and the other natural language observation. First, we hypothesized that when listening context permits the preactivation of words, sensorimotor brain regions associated speech production will be engaged to activate *specific* words *before* those words are heard compared to less predictive contexts. For example, speech production regions would be more activate before ‘frogs’ in the ‘pond’/’croaking’ sentence compared to ‘frogs’ in the sentence ‘The woman talked about the frogs’. Second, we hypothesized that, if the result of using listening context is a reduction in processing demands, AC will be less active in predictive compared to less predictive listening contexts at the time the more predictable word is spoken.

## fMRI study one

To test these hypotheses, 12 participants (N=7 females, 4 males, 1 unreported; average age of those reporting = 29.28, SD = 6.18) underwent fMRI while listening to experimentally manipulated sentences that sometimes contained disfluencies. Disfluencies included both unfilled pauses (with no sound) and pauses filled with the occasional ‘um’. To explain these pauses, participants were told that sentences were recordings of an actor at various stages in the process of memorizing his lines and, therefore, spanned a range of fluency from imperfect to well rehearsed. Disfluencies were included so that brain responses to sentence context could be statistically separated from the response to words associated with that context.

There were three sentence types, filler sentences and high- and low-predictability sentences. Unanalyzed filler sentences contained either no disfluencies or one unfilled or filled pause at the beginning or middle of the sentence. Analyzed high- and low-predictability sentences each consisted of a sentence frame (with an ‘um’ following the first word), a subsequent pause of variable duration with at least one ‘um’ and then a final word (Figure 2a). The final words were acoustically the same in both high- and low-predictability sentences. Predictability of these words was based on the sentence frame content, established in another experiment, and were otherwise balanced for intelligibility, phonetic content and length^27^. We used natural or ‘passive’ listening in this and study two because our hypotheses involve sensorimotor systems and an overt task would require confounding motor responses. Brain imaging data were analyzed using a deconvolution/regression model and hypotheses were tested by statistical contrasts of resulting activity during the filled pauses and final words as described next.

**Figure 2.**
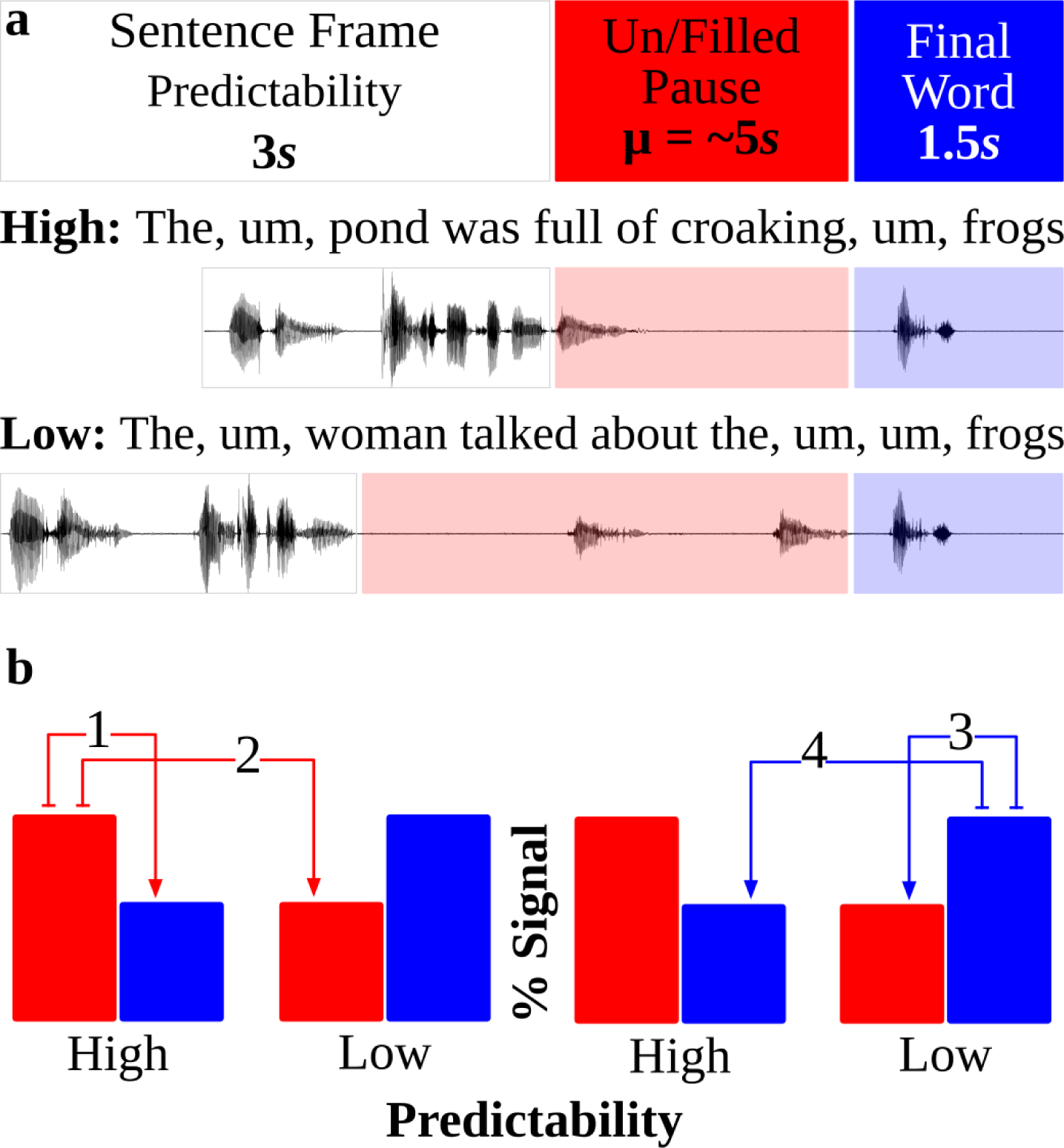
Design and analysis logic of fMRI study one. a) Participants listened to randomly presented sentences. Analyzed sentences had sentence frames, serving as context, making the final words either high- or low-predictability. Sentence frames were separated from the final word by both filled (‘um’) and unfilled pauses of variable duration to permit hypothesis testing. b) Graphical depiction of hypothesized results. The y-axis is percent (%) signal change of brain activity in auditory and speech production related regions. Numbers refer to statistical contrasts described in the text pertaining to the filled pauses (1/2; red) and final words (3/4; blue).

### Activity before final words

Examining the time period before the final words addresses the question of what, if any, brain regions are involved in exploiting words preactivated by listening context. If processes and regions associated with speech production are ‘reused’ prior to the final word, both of the following contrasts should activate speech production related regions (Figure 2b):

1. High-Predictability Filled Pause > Low-Predictability Filled Pause
2. High-Predictability Filled Pause > High-Predictability Final Word

The rationale by the hypothesis-and-test model (Figure 1) is as follows: 1) Specific words are more likely to be preactivated during the sentence frames and used by the speech production network during the filled pause for high-predictability sentences than low-predictability sentences. 2) If the result of engaging the speech production network during the high-predictability filled pauses is a decrease in processing demands, there should be less need for these regions during the high-predictability final words.

The intersection of contrasts 1) and 2) showed activity in sensorimotor regions typically thought to be involved in speech perception and production^22^. These include bilateral posterior superior temporal cortices, inferior parietal lobule (IPL), the pars opercularis (POp; ‘Broca’s area’), supplementary motor area (SMA), pre- and primary motor cortices and the cerebellum (Figure 3 left column and white outline; Table S1). To examine the perceptual, motor and cognitive processes engaged during the filled pauses, we correlated the spatial pattern of activity from this time window with 12 meta-analyses of neuroimaging data, returning a single correlation coefficient per pair^28,29^. The assumption is that if the spatial pattern during high predictability filled pauses resemble, e.g., the pattern from studies of oral/facial movements, that some element of the latter process is engaged at this time period. Indeed, though all the participants ostensibly heard during pauses was ‘um’, brain activity for the high-predictability filled pause was more correlated with activity associated auditory and word processing and oral/facial and speech production movements than activity from the low-predictability filled pauses (Figure 4a).

**Figure 3.**
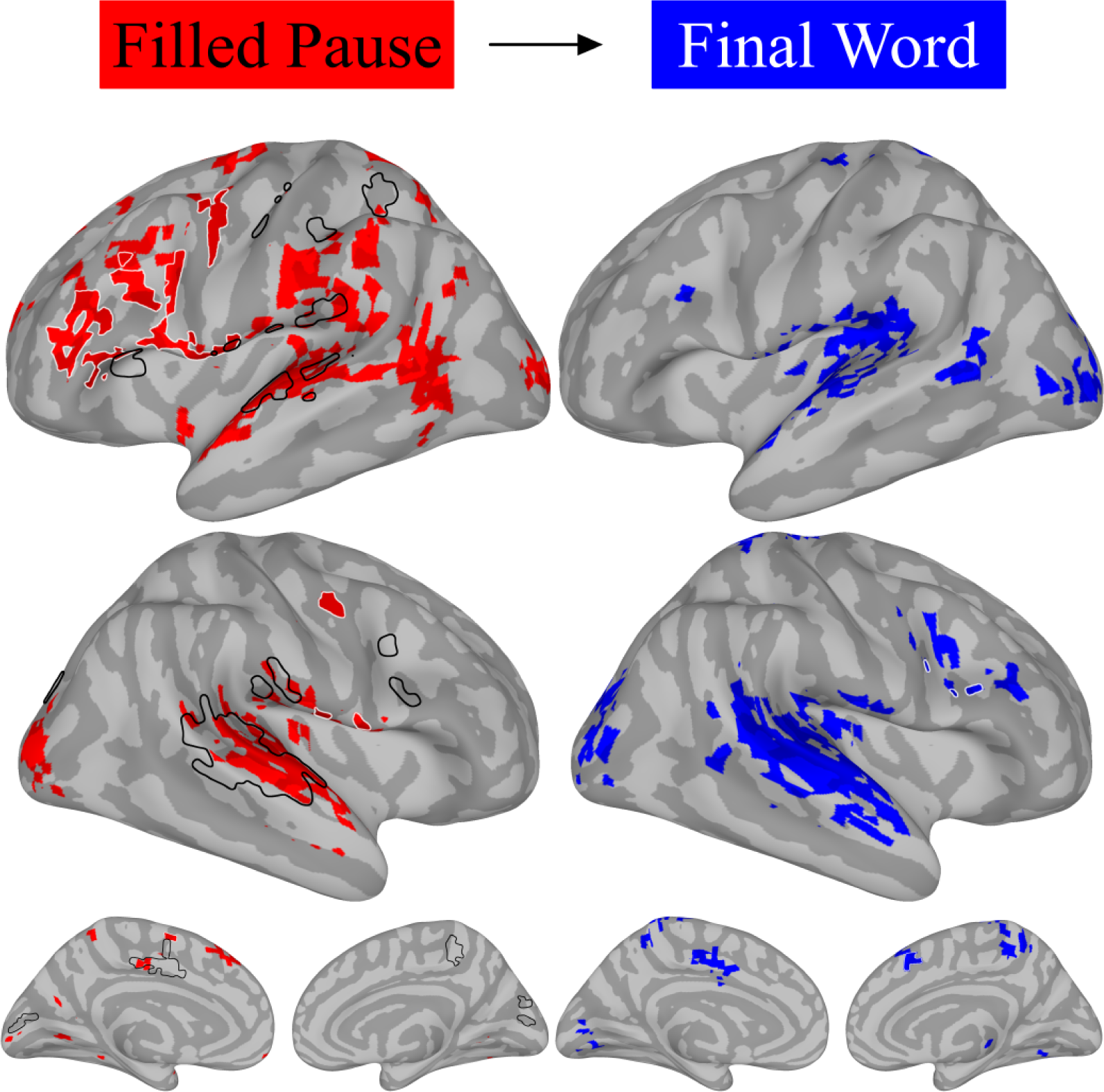
fMRI study one activity by predictability. Left column brains show greater activity during filled pauses following high-predictability sentence frames (1/2 in Figure 2b). The right column shows greater activity during final words following low-predictability sentence frames and filled pauses (and implies the converse, less activity for high-predictability final words; 3/4 in Figure 2b). Black outlines are regions supporting accurate classification by a support vector machine of high- but not low-predictability filled pauses as words when trained on final word activity. White outlines are regions overlapping those involved in making oral/facial movements (from Figure 1).

**Figure 4.**
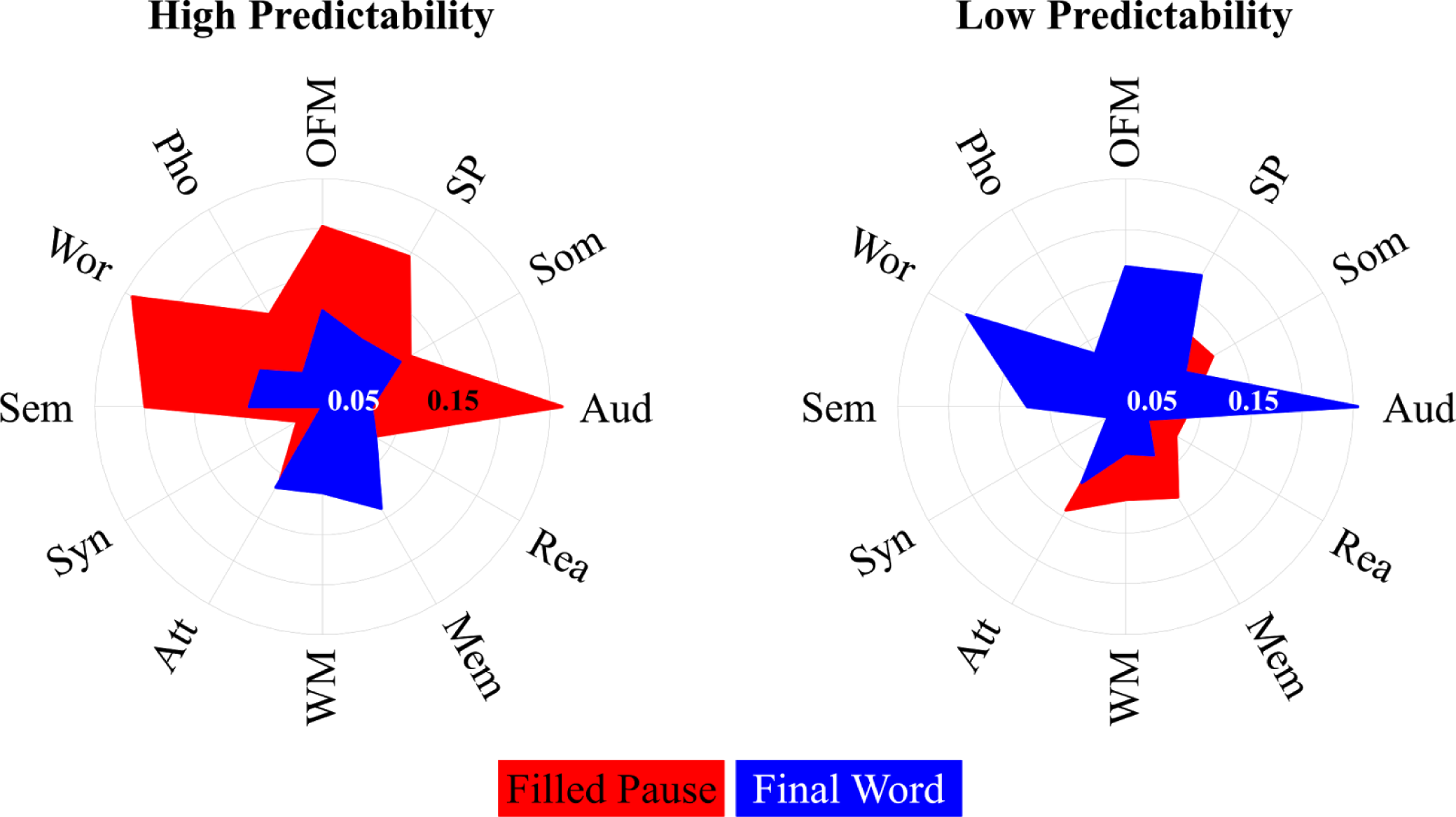
Correlation of study one results with fMRI meta-analyses. Radial plots of the correlation coefficients from the correlations of the spatial patterns of activity from high- and low-predictability filled pauses (red) and final words (blue) with the patterns from 12 fMRI meta-analyses corresponding to labels. Abbreviations: Aud = Auditory; Som = Somatosensory; SP = Speech Production; OFM = Oral/Facial Movement (shown in Figures 1 and 3); Pho = Phonology; Wor = Words; Sem = Semantics; Syn = Syntax; Ate = Attention; WM = Working memory; Mem = Memory; Rea = Reasoning.

### Activity during final words

Examining the time period following the filled pauses addresses the question of what the neurobiological consequences of using words preactivated by listening context are. If speech production regions are ‘reused’ in a predictive manner to constrain interpretation of acoustic patterns arriving in AC, both of the following contrasts should evince greater brain activity (Figure 2b):

3) Low-Predictability Final Word > High-Predictability Final Word
4) Low-Predictability Final Word > Low-Predictability Filled Pause

The rationale as inferred from the hypothesis-and-test model (Figure 1) is as follows: 3) It would not be until the low-predictability final words begin that AC could start processing those words. In contrast, prediction would have already been engaged during the high-predictability filled pauses. This would result in a reduction in processing demands during the high-predictability final words, associated with less activity. 4) The low-predictability final words would also produce more activity than the low-predictability filled pauses. This is because the low-predictability sentence frame could not be used to engage speech production related processes and, thus, should result in less activity during the low-predictability filled pauses.

The intersection of contrasts 3) and 4) showed a large amount of activity primarily in AC, lacking or having a significant reduction of activity in many of the regions in 1) and 2) that might be described as being related to speech production. (Figure 3 right column; Table S2). To determine the processing roles most associated with the distribution of activity during final words, we again correlated the spatial pattern of activity with 12 neuroimaging meta-analyses. The high predictability final words were less robustly correlated with oral/facial and speech production movements compared to the high-predictability pauses (Figure 4b).

We conducted two additional analyses. First, we tested the hypothesis that *specific* words are preactivated and selected from high-predictability sentence contexts. To do this, we trained a support vector machine (SVM, ^30^) on the high- and low-predictability final words. We then tested its accuracy at classifying activity during the high- and low-predictability filled pauses (the time windows in which participants heard only ‘um’s or nothing). The logic was that, if words are preactivated and selected from the high-predictability sentence context, a classifier should ‘confuse’ activity *before* the word in the high-predictability case with the activity for the word heard in the low-predictability case (i.e., it should classify high-predictability filled pauses as low-predictability final words). It should not confuse activity from the high-predictability filled pauses with the high-predictability final words because, by our model, those words do not need to be processed fully.

Before testing the specificity hypothesis, we first determined if a classifier could distinguish between the activity patterns associated with high- and low-predictability words. We trained a SVM on half of the high- and low-predictability final words and tested it on the other half and vice versa. Indeed, high- and low-predictability final words could be distinguished with classification accuracies being 57.01% and 58.90% respectively (with no difference in accuracy; *M* difference = −1.89%; t(23) = −0.93, p = 0.36). To test the specificity hypothesis, we trained an SVM on all of the high- and low-predictability final words and tested it on the high- and low-predictability filled pauses. Indeed, the the high-predictability filled pauses were ‘accurately’ classified as low-predictability final words 60.99% of the time compared to 39.77% for the low-predictability filled pauses (*M* difference = 21.22%; t(11)=3.30, p=0.007). Activity supporting this classification was determined by a group level one sample t-test using the weighted linear support vectors from each participant. Activity was in AC, primarily posterior superior temporal cortex (STp), parietal, POp, insula, SMA and motor and somatosensory cortices (Figure 3 black outline; Table S3).

Second, we used exploratory network analysis to test the hypothesis that preactivated words serve as ‘hypotheses’ that are ‘tested’ through feedback (or efference copy) to AC in high-predictability contexts. Specifically, we used bivariate autoregressive modeling^31^ to find the feedforward and feedback connectivity between each voxel in the brain with every other voxel in the brain separately for the high- and low-predictability filled pause and final word time period. We then counted the number of significant feedforward and feedback connections associated with each voxel and used t-tests to determine if there was a significant difference in the number of feedforward or feedback connections between the high- and low-predictability sentences. The logic is that, if context is used to constrain processing in AC, these regions should be the recipient of a significantly greater number of feedback connections during high-predictability sentences. Indeed, when comparing the high- and low-predictability sentences, the low-predictability sentences resulted in significantly more feedforward connections throughout the brain (Figure 5 seed-to-target, blue; Table S4). In contrast, there was significantly stronger feedback connectivity for the high- compared to low-predictability sentences in primary AC, posterior superior temporal cortices, IPL, somatosensory and premotor regions (Figure 5 target-to-seed, red; Table S4).

**Figure 5.**
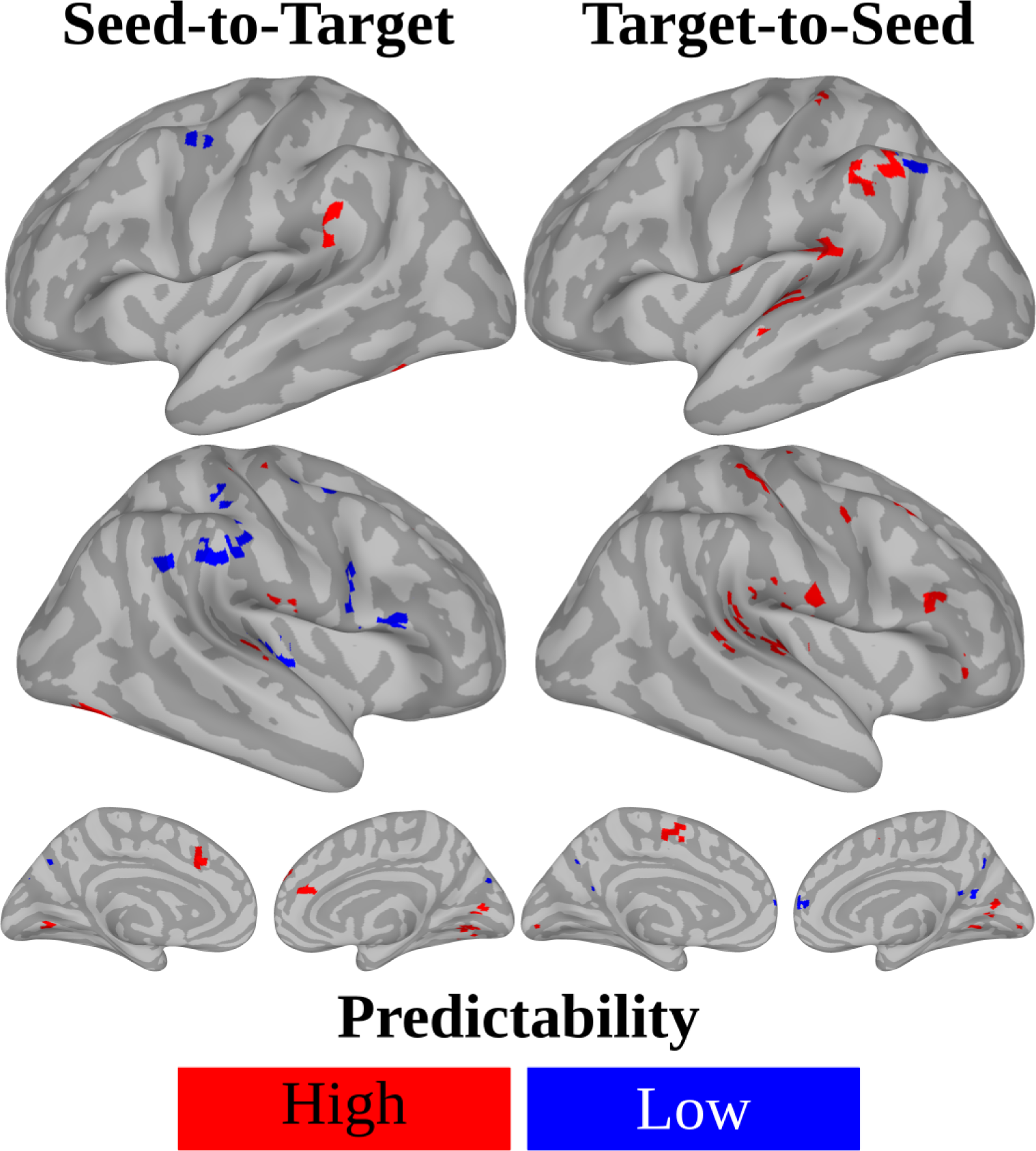
fMRI study one network connectivity by predictability. Brain images show a significantly greater number of seed-to-target (‘feedforward’; left) or target-to-seed (‘feedback’; right) connections for high- (red) and low-predictability (blue) filled pauses and final words.

To summarize, results cohere well with the proposed hypothesis-and-test model (Figure 1) and two overarching hypotheses. Affirming hypothesis one, results suggest that sensorimotor regions involved in speech production are engaged more in high compared to less predictable listening contexts *before* the onset of the words made predictable by that context (Figure 3 left column white outline). Though participants heard only meaningless sounds during this time window, activity patterns in part resemble activity that occurs when people listen to and produce meaningful words (Figure 4). Furthermore, a machine learning approach showed that those patterns resemble the *specific* words participants would hear next (Figure 3 left column black outline). Affirming hypothesis two, when final words are actually presented, activity in AC is significantly decreased for more compared to less predictive sentences contexts (Figure 3 right column). Even though participants were listening to the same final words in both cases, it seems as if little meaningful word processing was happening for the high-predictability final words (Figure 4). This reduction in speech processing in high-predictability contexts corresponds to AC becoming the stronger feedback recipient of influences from other brain regions (Figure 5).

## fMRI study two

Study one stimuli were unnatural in a number of ways that lead us to test the same two overarching hypothesis using a real-world stimulus. First, context is not only verbal and sentential as in study one: natural context includes discourse speech-associated mouth movements and co-speech gestures. These movements are used by the brain in the process of speech perception and language comprehension^11,20^’^32^’^33^. Second, though filled pauses are more characteristic of natural speech than completely fluent sentences, the duration of these pauses is likely shorter and more linguistically variable than in study one. This delay might have encouraged participants to use an overt guessing strategy, or biased them toward anticipatory lexical processing. Participants did not verbally report trying to predict the final word of sentences nor did they know what the experiment was about in a post scan questionnaire (perhaps because of the cover story and filler sentences). Nor is there reason to expect that implicit, as compared to more overt, prediction would result in qualitatively different activity patterns. Nonetheless, study one stimuli lead to the question of whether observed activity patterns will generalize to stimuli with other naturally varying contexts with typical delays and disfluencies.

To extend results to more natural speech and variable forms of context, we had fourteen participants (6 females, 8 males; average age = 24.6, SD = 3.59) watch a television game show with natural audiovisual dialogue. To be able to test the two overarching hypotheses also tested in study one, we first conducted a separate web-based crowdsourced experiment to determine which words in the show were high- and low-predictability. Participants rated each word for how predictable it was based on what had come before it in the show. Resulting scores ranged from 4.70 - 99.58 on a 100 point scale and the median/mean predictability score was 62.00/62.31. We then median split all the words into two categories, high- (median to maximum) and low-predictability (minimum to median) words.

In order to determine the distributions and differences in brain activity for these word categories, a new method to analyze neuroimaging data from natural continuous stimuli was needed. First, we use a model free multivariate approach, spatial and temporal independent component analysis (stICA)^34^’^35^, to separate the brain data into a set of networks and corresponding timecourses for each participant. Each of these unmixed networks is theoretically involved in processing some specific aspect of the show, including the high- and low-predictability words. Note that this unmixing cannot be done with traditional analysis approaches unless one has a priori knowledge about what in the show drove every response in the resulting data (an impossibility that is typically resolved by using reductionist and constructed stimuli like those in experiment one).

To identify which of the resulting networks is associated with processing high- and low-predictability words, a ‘reverse correlation’ approach called ‘turnpoints analysis’ is employed^32,36,37^. In particular, an stICA network is said to process high- or low-predictability words if those words occur more when the brain response in each associated network timecourse is rising than when falling. Conversely, a component is said not to process those words if they occur more when the response is falling than rising. This is determined by counting the number of times words aligned with rising (called peaks) or falling responses (called valleys) and comparing resulting counts using a chi-square test.

This comparison is done on each network timecourse at zero lag and at increments of 1.5 seconds for six lags. This nine second time window is chosen because it is assumed that stICA network timecourses follow principles of the ‘canonical’ hemodynamic response function (HRF) used in the analysis of most fMRI data. In particular, the HRF for a ~500ms word in isolation (without any context) would be about nine seconds long. The HRF for this word would not begin to rise until about two seconds after its onset. It would then rise and reach a peak about two seconds later and decrease and return to baseline about five seconds after this. All resulting stICA networks associated with high- and low-predictability words at peaks > valleys and valleys > peaks are separately combined by summation, resulting in four network maps per lag per participant.

Finally, group analyses were done using SVM classifiers and analysis of variance (ANOVA). SVMs were used to determine if network maps can be used to distinguish between whether participants were listening to high- or low predictability words and, if so, what brain regions support those classifications. SVMs were trained on half of the participants high- and low-predictability network maps and tested on the other half of the participants at each lag. To support the validity of this approach and extend results, we also conducted more traditional group analysis by univariate three-way repeated measures ANOVA of network maps, with predictability, lags and participants as factors.

### Activity before words

To review, hypothesis one maintains that activity in sensorimotor brain regions occurs before word onset for high- but not low-predictability words, corresponding to the selection, sequencing and prediction of forthcoming speech sounds from context. This hypothesis was tested with the lag spanning 0-1.5 seconds, corresponding to the onset of words. By typical assumptions about the HRF reviewed above, accurate classification and this time window suggests that the classifiers can predict which category of word was to be heard *before* it could have been heard. Indeed, SVMs could accurately distinguish high- and low-predictability words at 0-1.5 seconds (Figure 6 bottom left). Classification was based on a distributed set of sensorimotor networks associated with high-predictability words. This included two large bilateral STp clusters in addition to parietal, SMA and premotor cortices and the caudate and cerebellum. At valleys classification was based in pre- and primary motor cortices (Figure 6; Table S5). Based on standard assumptions about the HRF, activity in valleys at this lag suggests that high-predictability word responses in the latter regions had peaked and were in the decreasing amplitude phase of the HRF. That is, processing in these regions began before peaks in STp cortices. In contrast, classification associated with low-predictability words was mostly limited to small clusters in pre- and primary motor cortices (Figure 6; Table S5). These results were largely confirmed by second order contrasts for predictability following ANOVAs at this time window with the exception that there was no significant activity associated with low-predictability words (Figure S1).

**Figure 6.**
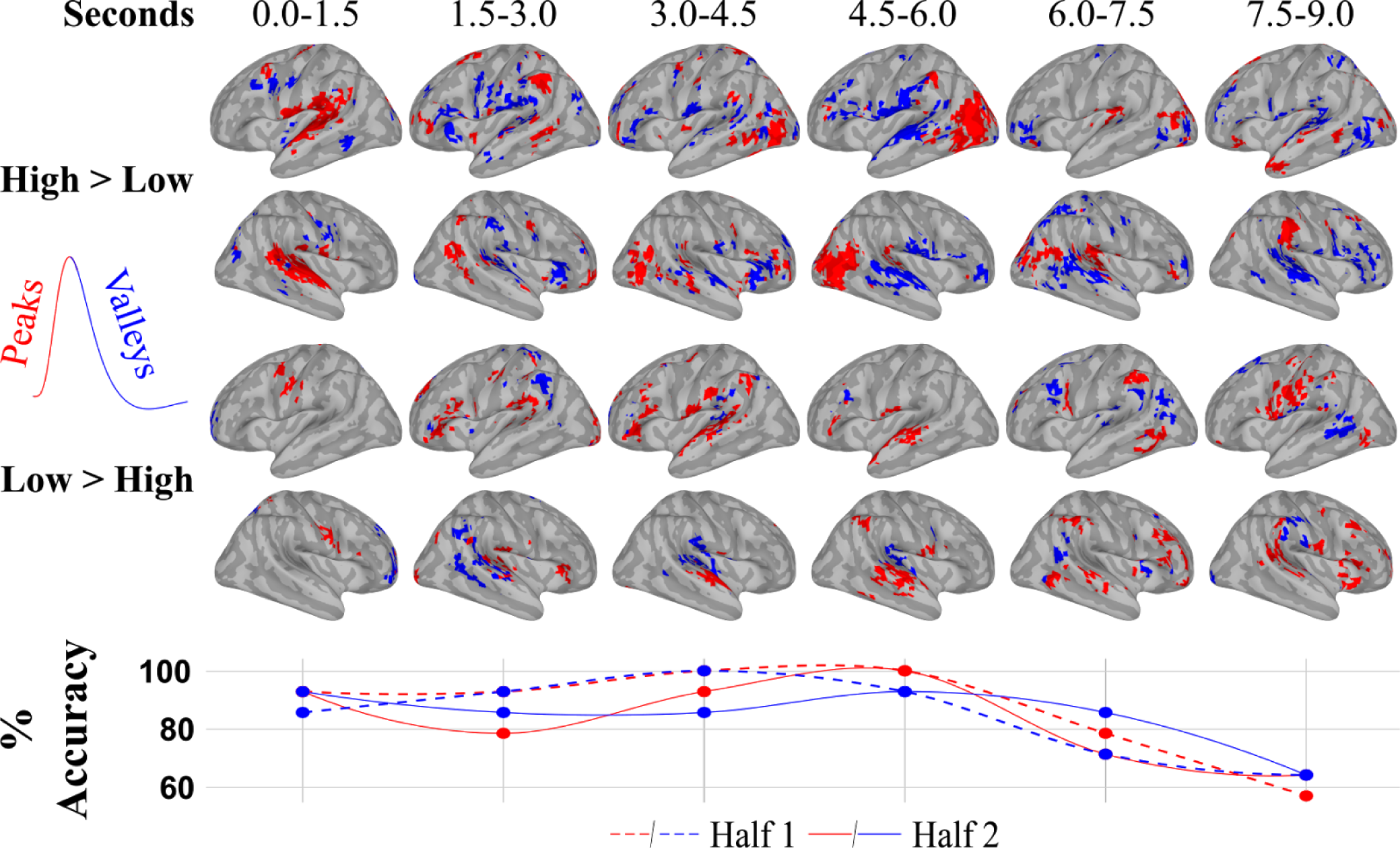
fmri study two word predictability turnpoints analysis. Turnpoints analysis as described in the text was done on each participant to find independent component networks corresponding to high- and low-predictability words in a television game show. Support vector machine (SVM) classifiers were trained to distinguish those word categories on half the participants and tested on the other half. SVMs were done at six time windows (‘Seconds’; top) for networks corresponding to words when brain responses were rising (‘Peaks’; red) and falling (‘Valleys’; blue). Classification accuracy is given in the bottom row (min = 57.14; *M*= 84.82; max = 100.00).

### Activity during words

To review, the second hypothesis maintains that there is a reduction in activity in AC for high-compared to low-predictability words corresponding to a processing savings following accurate word prediction. Indeed, at lags of 3-9 seconds, the ability of the SVM to accurately determine whether participants had been listening to high- and low-predictability words was based on a distributed set of regions mostly associated with a decrease in brain responses for high-predictability words and an increase for low-predictability words (Figure 6; Table S3). Decreases in responses for high-predictability words were in sensorimotor regions, including bilateral STp, IPL, POP, SMA, insula and pre- and primary motor cortices. There was also a large increase in response in visual cortices. Increases in responses for low-predictability words was mostly in bilateral STp. Frontal and parietal activity patterns for low-predictability words was less extensive than those associated with high-predictability words. The general pattern of activity for low-predictability words is much like what would be expected of a typical HRF for a word in isolation as described above: little to no activity until two seconds and a peak at four seconds, followed by a slow decrease in activity. Again, this pattern of results were largely confirmed by second order contrasts in predictability following ANOVA at these time windows (Figure S1; Table S5). The interaction of predictability and time further confirm the overall pattern of activity before and after high- and low-predictability words as involving mostly sensorimotor regions (Figure S1 white outline; Table S6).

To summarize, study two affirms that study one results generalize to real-world stimuli with multisensory context and natural speech timing. Supporting hypothesis one, sensorimotor networks associated with speech production were engaged prior to high-predictability word onsets. In contrast, there was little or no activity in these networks for low-predictability words at this time window (Figure 6 0.0-1.5 seconds). This pattern of activity prior to word onset also suggests that processing of high-predictability words involves feedback from frontal production related to auditory regions. Supporting the second hypothesis, there was a large decrease in brain activity associated with high-predictability words throughout sensorimotor brain networks following word onset. There was a concomitant increase of activity in visual networks that, speculatively, might correspond to increased visual sampling of the video because of freed up processing demands associated with having already predicted the subsequent word. In contrast, low-predictability words seem to be primarily associated with positive activity in mostly auditory networks that peak at a later time (Figure 6 1.5-9.0 seconds).

Thus, in two studies, results support an hypothesis-and-test model of speech perception (Figure 1). By this model, the speech production system is ‘reused’ so that words activated or primed by listening context through probabilistic associations can be selected, sequenced and used to predict forthcoming speech sounds. Indeed, both studies showed that temporally evolving sensorimotor brain networks involved in speech perception and production are active in advance of words when context can be used to predict those words. Patterns of activity in those networks are associated with specific language processes and even specific words, namely those that are to be heard next. Furthermore, AC became the feedback recipient of information from the rest of the brain generally and frontal speech production related regions more specifically in more predictive contexts. Thus, results confirm that specific unheard ‘hypotheses’ are generated by the brain from contextual associations. The model then proposes that these hypotheses are ‘tested’ by comparing them to incoming information to constrain interpretation of variant acoustic patterns. If this is true, accurate predictions (as would derive from high predictive contexts) result in a reduction in processing demands and associated brain activity. Indeed, in both studies, at the time the to be heard words are actually heard, there is a subsequent and large reduction in brain activity in high-predictability contexts. Thus, results support the contention of this model that, when it is available, context is used as a constraint to achieve perceptual constancy in the process of speech perception. As context is abundant during natural language use, results suggest that a great deal of the speech we hear is an active knowledge based construction, built more from our associations with context than from the incoming auditory information itself.

Results have implications not only for how we perceive speech but also for models of the organization of language and the brain. Most classical and contemporary models of the neurobiology of language are static, with speech perception and language comprehension proposed to occur in a fixed region (like ‘Wernicke’s area’)^38^ or fixed sets of connected regions (like the ‘dorsal’ and ‘ventral stream’)^39^’^40^. The results of our two studies, however, show that the general patterns of activity and network connectivity for words, even for exactly the same words, are remarkably different as a function of the context encountered by listeners. If different patterns of activity are associated with different contexts, and contexts are always changing during real-world language use, it implies that the organization of language and the brain is much less static and more dynamic than existing models can accommodate. Indeed, studies of the network organization supporting resting states, auditory tone processing and natural language all suggest that language and the brain is dynamic and distributed throughout the brain^22,32,41–43^. These studies and our results thus suggest that more research pertaining to how the brain uses context is needed in order to build a more complete and accurate model of the neurobiology of language^12^. A model that is more predictive of actual language behavior^12,44^ might help us to improve the generally poor outcomes in aphasia therapy^45^, a pressing concern given that aphasia results in the worst health related quality of life^46^.

## Methods

Two functional magnetic resonance imaging (fMRI) studies were conducted. Study one used experimental manipulation of stimuli. Study two involved natural observation of brain activity from watching a television game show.

### fMRI study one

#### Participants

There were 12 participants (7 females, 4 males, 1 unreported; average age of those reporting = 29.28, SD = 6.18). Each was a native English speaker, right-handed as determined by the Edinburgh Handedness Inventory^47^ with normal or corrected to normal hearing and vision and no history of neurological or psychiatric illness. The study was approved by the Institutional Review Board (IRB) of Weill-Cornell Medical College and participants provided written informed consent.

#### Stimulus and task

Participants listened to 40 high- and 40 low-predictability sentences and 43 filler sentences, as described in the manuscript. Stimuli were presented in a randomized rapid event related design, separated by an intertrial interval of variable duration. To be able to analyze activity associated with the filled pauses and final words, optimal stimulus randomizations were created using the programs ‘RSFgen’ and ‘3dDeconvolve’ from the AFNI program set^48^. Specifically, we ran one million iterations of ‘RSFgen’ to generate randomized stimulus functions using the Markov chain option to specify stimulus transition probabilities. The latter was needed because sentence frames must be followed by filled pauses and then final words. Randomizations were evaluated for efficacy using the the regression/deconvolution program ‘3dDeconvolve’^49^ in AFNI and the four most efficient designs were selected (one for each run).

Each of the recorded sentences was then reconstructed using the timings specified by these designs. This required splicing in prerecorded and natural sounding ‘um’ strings and room noise present at the time of the original recordings. Sentences were recorded in a room with some audible noise (the hum of a computer fan) so that there would be no audible evidence of splicing (namely clicks). After editing, sentence frames all occurred in a three second window. There were no significant differences in the resulting analyzed high- (*M=* 5.14, *SD =* 3.89) and low-predictability (*M=* 5.25, *SD =* 4.05) filled pause durations (*t*(39) =0.13, *p* = 0.89). There were also no differences in the duration and acoustic properties of the analyzed high- and low-predictability final words because they were the same word from the low-predictability version of the original sentence pairs. All final words occurred in a 1.5 second window.

Following scanning participants were asked questions about the stimuli to assess whether they attended during scanning. In particular, they read 24 sentences without the final word, 12 they had been presented in the scanner and 12 the had not been presented. They were asked 1) to complete the sentence 2) whether they had heard the sentence in the scanner and 3) their confidence that they had heard the sentence. Four participants were not given the questionnaire. Overall, the remaining eight participants were 61.77% confident in their answers. They were 75.20% accurate at identifying sentences that were actually presented in the scanner and 73.44% accurate at identifying sentences that were not actually presented in the scanner.

#### Imaging parameters

Brain imaging was performed at 3 Tesla (GE Medical Systems, Milwaukee, WI). A volumetric MPRAGE sequence was used to acquire anatomical images on which landmarks could be found and functional activation maps could be superimposed (Voxel size = 1.5×0.9×0.9 mm; Sagittal Slices = 120; FoV = 24). Functional imaging used an EPI sequence sensitive to BOLD contrast (Voxel size = 3.45 × 3.45 × 5; Axial Slices: 25; FoV = 220; Base resolution = 64; Time of Repetition, TR = 1500; TE = 30; Flip Angle = 75). There were four consecutive functional runs, each lasting six minutes and 21 seconds. Each run began with a 10.5 second silent period to allow magnetization to reach a steady state and these images were discarded.

#### Preprocessing

Unless otherwise noted, preprocessing was done with AFNI software^48^. Anatomical images were corrected for intensity non-uniformity, skull stripped^50^, non-linearly registered to an MNI template and inflated to surface based representations using Freesurfer software^51^. Freesurfer was used to create an average anatomical image from all participants from both studies one and two. All results were displayed on this average surface using the SUMA component of AFNI^52^

Functional images from the four runs were spatially registered in 3D space by Fourier transformation. A correction was applied for slice timing differences and spikes (signal intensities greater than 2.5 standard deviations from the mean) were removed. We then corrected each run for head movement by registration to the mean of of the middle run and aligned the results to the MNI template aligned anatomical images. We masked each run to remove voxels outside of the brain and created two sets of timeseries to be used in further analysis. In the first set, we blurred each run to a smoothness of 6 mm^53^ and normalized each run (by making the sum-of-squares equal to one). In the second set, we linearly, quadratically and cubically detrended each run, normalized and concatenated them. We submitted the latter to independent components analysis (ICA) to locate artifacts in the data^54^. In particular, resulting components were automatically labelled as not artifacts, possible artifacts and artifacts using SOCK^55^. Each was reviewed by hand for accuracy. There were an average of 207.33 components and 157.17 artifactual components (75.81%) across participants. The independent component time course associated with each of these was removed from the timeseries at each voxel using linear least squares. Finally, we blurred the resulting timeseries to a smoothness of 6 mm.

#### Regression/deconvolution analyses

The four blurred and normalized timeseries (set one) were used to estimate the hemodynamic response for the high- and low-predictability filled pauses and final words in each voxel using ‘3dDeconvolve’ from AFNI^49^. In particular, we used a deconvolution approach to produce an unbiased estimate of the time course for these stimuli following stimulus onset with piecewise cubic-spline basis functions. These were separated by 1.5 second intervals and covered a 15 second time window. Thus, each condition produced four timeseries, each with 12 coefficients (or timepoints) per voxel. In addition, the model included the ICA based artifact time courses as regressors, and a regressor each for the mean signal, linear, quadratic and cubic trends.

#### Overlap analysis

To test hypotheses described in the manuscript, each of the resulting four estimates of the hemodynamic response was used to calculate a paired t-test at each timepoint following stimulus onset. Specifically, we performed paired t-tests for high- vs. low-predictability filled pauses; high-predictability filled pauses vs. high-predictability final words; low- vs. high-predictability final words and; low-predictability final words vs. low-predictability filled pauses. Each timepoint for contrasts 1-4 described in the manuscript was thresholded. To correct for multiple comparisons, we used an individual voxel threshold for each overlapping image that equaled p <.005 when combined and a cluster size threshold of 10.5 voxels. These were determined by Monte Carlo simulation to correspond to a corrected alpha value of .05 using ‘3dClustSim’ (note that we use a version of this program that fixes a previous bug). Finally, we collapsed over time by summation.

#### Classifier analysis

To discover if words are activated during the high-predictability filled pauses, we conducted the classification analyses described in the manuscript. In particular, the deconvolution analysis was rerun but by splitting the high- and low-predictability filled pauses and final words into even and odd stimulus presentation onsets. This resulted in eight timeseries. We trained a support vector machine (SVM) classifier on the even high- and low-predictability final word timeseries and tested it using the odd high- and low-predictability final word timeseries and vice versa. We similarly trained a SVM on the high- and low-predictability final word timeseries and tested it with the high- and low-predictability filled pause timeseries. Analyses used ‘3dSVM’^30^. Group analysis was done using a one sample t-test as described. Thresholds were determined using a mask of the overlap of high- and low-predictability filled pauses versus baseline, an individual voxel p-value of .05 and a cluster size of 14, resulting in a corrected alpha level of .05 as determined by ‘3dClustSim’.

#### Network analysis

We conducted network analysis by first creating a mask that was the sum of contrasts 1)-4) as described in the manuscript and their converse to reduce the total number of computations. We then did exploratory bivariate autoregressive modeling with two lags at each of the 10,113 voxel in this mask. Resulting path estimates (and associated t-statistics) give a measure of the potential connectivity between each seed voxel and the rest of the voxels in the brain (seed-to-targets) and from the rest of the brain to the seed voxel (targets-to-seed). We minimally thresholded each image at p = .05 uncorrected with a cluster size of five voxels and counted the number of active voxels at each voxel across all 10,113 resulting maps. We used paired t-tests to contrast high- vs low-predictability filled pauses for each lag and for both seed-to-targets and targets-to-seed separately. Thresholds were determined using the mask used to constrain computations, an individual voxel p-value of .005 and a cluster size of 10, resulting in a corrected alpha level of .05 as determined by ‘3dClustSim’. Results as displayed in Figure 5 are collapsed over lag. of the overlap of high- and low-predictability filled pauses versus baseline

#### Neuroimaging meta-analyses

We correlated the spatial patterns of results form study one with neuroimaging meta-analyses conducted using the BrainMap database (http://brainmap.org/ySpecifically, we queried that database for experiments meeting a set of common metadata criteria and sets of criteria specific to 12 perceptual, motor and cognitive processes. These queries returned x/y/z stereotaxic coordinate space “locations”, that is, centres of mass or peaks of functional brain activity reported in neuroimaging papers^56–58^. Locations that were originally published in the Talairach coordinate space were converted to Montreal Neurological Institute (MNI) space^59,60^. Then Activation Likelihood Estimation (ALE) meta-analyses were done by modelling each MNI location as a three-dimensional probability distribution and quantitatively assessing their convergence across experiments. Significance was assessed by ten thousand permutations of above-chance clustering between experiments^61–64^. All resulting ALE maps were false discovery rate (FDR) corrected for multiple comparisons to p < 0.05 and further protected by using a minimum cluster size of 84 mm^3^ (10.5 voxels).

The BrainMap database was searched (in March, 2016) for a set of common criteria associated with all 12 meta-analyses. In particular, experiments contributing to analyses included only “normal” participants who were right handed and older than seventeen and locations must be activations only. These common search criteria were combined with searches for for 1) audition and 2) somesthesis in the behavioral domain of perception. They were combined with searches for 3) attention; 4) memory; 5) phonology; 6) reasoning 7) semantics; 8) syntax; and 9) working memory in the behavioral domain of cognition. They were combined with a search for stimulus types that were auditory 10) words. They were combined with a search for 11) speech production in the behavioral domain of action. Finally, they were combined with a search for 12) oral facial movements that variously involved overt oral/facial responses of breath-hold, drink, smile, swallow or the paradigm classes of breath holding, chewing/swallowing, eating/drinking, swallowing and taste without auditory stimuli. It was not the intention that these meta-analyses represent independent constructs but, rather to examine the correlation of our experimental results with a wide range of studies examining various processes. We used ‘3ddot’ in AFNI to correlate the spatial distribution of activity from the overlap of contrasts 1) and 2) and their converse and the overlap of 3) and 4) and their converse (Figure 3) with these 12 meta-analyses (resulting in 48 correlations) to provide a description of the processes engaged during the high-predictability filled pauses and final words.

### fMRI study two

#### Participants

There were 14 participants (6 females, 8 male; average age = 24.6, SD = 3.59 years). All other participant criteria were the same as in study one. None of the study two participants took part in study one. IRB approval for scanning was the same as for study one whereas approval for the predictability rating study was from Hamilton College.

#### Stimulus and task

Participants listened to and watched 32 minutes and 24 seconds of an episode of a television game show (“Are You Smarter Than A 5th Grader”; Season 2, Episode 24, Aired 2/7/08). The episode was edited down to be approximately 30 minutes without decreasing intelligibility. The show was further divided into six segments so that we could check on participants and give them breaks. This show was chosen because it had a number of desirable properties including: 1) Natural dialogue between the host, contestant, and, to a lesser extent, six peripheral individuals (i.e., no actors); 2) Naturally occurring temporal jittering of dialogue and contextual information, allowing stimulus features to be resolved using fMRI; and 3) A built-in behavioural assay to assess participants knowledge of the items or events discussed in the video and to determine whether participants were attending. Participants who completed this assay following the experiment (N=10) were on average 98% accurate and 82% confident in their answers when asked the contestant’s answers to 11 questions during the show. The contestant on the show answered all questions correctly except the last. Participants were 80% accurate when asked the contestant’s answer to the final question after having seen the show despite that only 30% indicated that they had known the answer to the question before viewing the show (z(9)=2.2473; p < 0.02).

#### Stimulus annotation and word predictability

Praat^65^ and ELAN software were used to annotate the onset and offset time of words in the TV show episode watched by participants. Following annotation, the words were resampled to 4 ms time steps so that events could be aligned to fMRI timeseries generated by the fMRI analyses and compared to EEG data whose sampling rate was 250 hz (not discussed in this manuscript). Resampled word annotations were entered as a table into a MySQL database (http://www.mvsql.com/) along with metadata about how predictable each word was based on the context preceding it. These were determined in a separate experiment using Amazon’s Mechanical Turk (www.MTurk.com) and Qualtrics survey software (http://www.qualtrics.comy).

Specifically, six groups of native English speaking participants rated words in one of six transcripts (N = 291 participants; M = 48.50 and SD = 8.26 participants per transcript). Multiple groups were used to break the task up in the same way as the fMRI experiment (with six functional runs) and because the time required to rate more than one transcript would have been too great (each took about 1. 5 hours). None of the MTurk participants participated in the fMRI experiment. Participants rated each word in the transcript that they were assigned on a continuous Liket scale. The scale was labeled 0 to 100 in intervals of 10 and also included the words “Strongly Unpredicted” (at 0), “Unpredicted” (at 25), “Medium Predicted” (at 50) “Predicted” (at 75) and “Strongly Predicted” (at 100 on the scale). Participants were given instructions to indicate how predicted each word was based on what they had previously read and were provided examples of how to do so with the Likert scale. After reading and rating a word they advanced to the next word and were not able to see prior or upcoming words. If participants were not in the transcript one group, they read all the preceding transcripts in the correct temporal order before they started rating.

To get reliable estimates of word predictability from the MTurk groups, the original groups were narrowed down by discarding participants who consistently made extreme word ratings. This was done by making a box plot of the ratings for each word for all participants and marking individual participants as outliers if they rated any given word below Q1 - 1.5×IQR or above Q3 + 1.5×IQR. The number of outlying words were then summed for each of the participants. Those participants whose ratings produced more outliers than the median number of outliers were excluded from the estimate of word predictability. This left us with 119 or 40.89% of the the original number of participants (M = 19.83 and SD = 1.33 participants per transcript; 65 males; age M = 34.13 and SD = 11.44). Intraclass correlation coefficient (ICC) confirmed that trimmed groups were relatively consistent (ICC M = 0.57 with ICC range of 0.48 - 0.64). We then simply averaged the remaining participants ratings for each word. The resulting minimum rating was 4.70, Q1 was 41.25, the median was 62.00, the mean was 62.31, Q3 was 85.50 and the maximum rating was 99.58. We defined high-predictability word as those words from medium to maximum predictability and low-predictability words as those from the medium to lowest predictability.

#### Imaging parameters and preprocessing

There were six consecutive functional runs with durations, in minutes:seconds of 5:36, 5:45, 5:42, 5:42, 4:51 and 5:51. Each run began with a 10.5 second black screen that faded out to allow magnetization to reach a steady state and these images were discarded. All parameters and preprocessing steps were the same as in study one. With regard to the ICA based artifact discovery, there were an average of 237.29 components and an average of 177.79 artifactual components (74.93%) per participant.

#### Turnpoints analysis step one

To find independent brain networks without a priori hypotheses about where they are or what they are doing, each participant’s preprocessed and smoothed timeseries (with artifacts removed) was submitted to both spatial and temporal independent component analysis (stICA)^34^’^35^. Number of components were estimated using an automatic heuristic strategy^34^. The independent component (IC) time-course for each of the resulting components was then resampled to have a 4 ms time step (to match the stimulus annotations and sampling rate of an EEG study not discussed here) and entered as a table into the MySQL database containing the annotations.

#### Turnpoints analysis step two

To determine which networks from step one were involved in processing high- and low-predictability words, we performed a ‘turnpoints peak and valley analysis’^32^ on the IC time-courses from the stICAs. A ‘turnpoint’ is defined as the point at which the time series transitions from ‘peak’ or ‘valley’ extrema points. Three successive data values are required to define a turnpoint: xt-1, xt, xt+1. Across a time series with n observations of xt for t = 2, …, n-1, a turnpoint xt is a peak if xt-l< xt> xt+1 or a valley if xt-l> xt< xt+1. That is, a peak is an observation that is both preceded and followed by a value lower than itself and a valley is an observation that is both preceded and followed by a value higher than itself.

The R statistical package (http://www.r-proiect.orgA) was used to querie the MySQL database and retrieve the high- and low-predictability word timeseries and an IC timecourses. If a high- or low-predictability word occurred when the IC time-course was rising (i.e., the time period between a valley and a peak) and did not occur when the response was decaying (i.e., the time period between a peak and a valley), then that timecourse and the associated spatial map were said to function to process that word. This was determined statistically using χ2 tests on the counts of an annotated feature at peaks and valleys measured in units of time. The unit of time in this case was 1.5 seconds (the TR). For example, if 100 words occurred at peaks and 10 at valleys and each word was exactly 450 ms long the values entered into a one-way *χ*2 would be 30 and 3 TRs. The expected counts under the null hypothesis would be 16.5 and 16.5 and *χ*2 = 22; p < 2.6x10-06. These *χ*2 tests were done over an extended time window (i.e., 0-9 seconds) in steps of 500 ms rather than at a fixed time. This was done because 1) of predicted differences in the timing of processing between high- and low-predictability words, 2) cognitive processing is extended over time and 3) a canonical hemodynamic response for an 500 ms word would extend over a window of approximately this length. Significance for components was set at p < .005.

#### Turnpoints analysis step three

At the conclusion of the last step, all significant stICA components involved in processing high- and low-predictability words have been found for each participant in a moving time window. Each map is z-score transformed and combined by summation into one map for each the high- and low-predictability words, for each the peaks and valleys, for each participant, for each of three consecutive 500 ms time steps (to correspond to the 1.5 second TR), resulting in four maps per participant for each of the resulting six time steps. Next, SVMs^30^ were trained using the high-and low-predictability word images for peaks and valleys separately from half the participants. We then tested the SVM using the other half of the participants and vice versa. In total we trained and tested four SVMs at each time point. The first half of the participants at peaks had a min = 64.29; M = 83.33 and max = 100.00 classification accuracy. The second half of the participants at peaks had a min = 57.14; M = 86.91 and max = 100.00 classification accuracy. The first half of the participants at valleys had a min = 64.29; M = 84.52 and max = 92.86 classification accuracy. The second half of the participants at valleys had a min = 64.29; M = 84.53 and max = 100.00 classification accuracy. In addition to accuracy, the results include those regions that contributed most highly to the classification across the training group.

## Supplementary Information

Is linked to the online version of the paper at www.nature.com/nature.

## Acknowledgements

This work was supported by NIH-NICHD K99/R00 HD060307 - ‘Neurobiology of Speech Perception in Real-World Contexts’ granted to JIS. JIS would also like to acknowledge EPSRC EP/M026965/1 ‘New pathways to hearing: A multisensory noise reducing and palate based sensory substitution device for speech perception.’ We would also like to thank John Pellman for his work on finding reliable word predictability ratings and helping write that section.

## Author Contributions

JIS and JZ designed the studies and collected the data; JIS analyzed the data and wrote the manuscript; JIS/JZ discussed the results and commented on the manuscript.

## Author Information

Reprints and permissions information is available at www.nature.com/reprints. The authors declare that they have no competing financial interests. Correspondence and requests for materials should be addressed to JIS.

## Supplementary Information

### Table Captions

**Table S1.** Filled pause activity locations (Figure 3 red). Abbreviations: AG = Angular Gyrus; AP = Anterior/Posterior; BA = Brodmann Area; IS = Inferior/Superior; L = Left hemisphere; R = Right hemisphere; RL = Right/Left; SMA = supplementary motor area; SMG = Supramarginal Gyrus.

|  | Region |  |  | Centre of Mass |  |  | Minimum |  |  | Maximum |  |  |
| --- | --- | --- | --- | --- | --- | --- | --- | --- | --- | --- | --- | --- |
| *Voxels/mm<sup>3</sup>* |  | Anatomical | BA | RL | AP | IS | RL | AP | IS | RL | AP | IS |
| 479/30656 | L | Superior/Middle Temporal Gyrus | 22/TE 3 | -53 | -17 | 9 | -30 | 45 | -17 | -70 | -68 | 40 |
| 264/16896 | R | Superior Temporal Gyrus | 22/TE 3 | 58 | -16 | 4 | 75 | 9 | -21 | 35 | -40 | 24 |
| 67/4288 | R | Middle Occipital Gyrus | 18 | 31 | -88 | 2 | 43 | -72 | -9 | 19 | -96 | 20 |
| 67/4288 | L | Inferior Frontal Gyrus | 44 | -42 | 8 | 26 | -34 | 29 | 8 | -50 | -8 | 40 |
| 64/4096 | L | Superior Frontal Gyrus | 6 | -14 | -5 | 62 | 3 | 9 | 48 | -26 | -20 | 76 |
| 64/4096 | L | Precentral Gyrus | 6 | -47 | -5 | 53 | -34 | 9 | 32 | -62 | -16 | 72 |
| 45/2880 | L | Middle Occipital Gyrus | 18 | -24 | -93 | 1 | -10 | -84 | -9 | -38 | -104 | 8 |
| 38/2432 | L | Superior Frontal Gyrus | 8 | -8 | 36 | 50 | -2 | 49 | 44 | -18 | 21 | 64 |
| 30/1920 | R | Inferior Parietal Lobule/SMG |  | 37 | -34 | 29 | 47 | -24 | 20 | 23 | -40 | 36 |
| 29/1856 | R | Precentral Gyrus | 4/6 | 57 | -5 | 46 | 63 | 5 | 40 | 47 | -12 | 56 |
| 26/1664 | L | Middle Occipital Gyrus | 31 | -22 | -62 | 25 | -18 | -56 | 16 | -30 | -68 | 40 |
| 19/1216 | L | Inferior Parietal Lobule/SMG | 40 | -55 | -43 | 38 | -50 | -36 | 32 | -62 | -48 | 48 |
| 17/1088 | L | Superior Parietal Lobule | 7A | -26 | -52 | 49 | -22 | -40 | 40 | -34 | -64 | 56 |
| 17/1088 | L | Superior Frontal Gyrus | 6 | -21 | 28 | 55 | -14 | 33 | 48 | -26 | 25 | 64 |
| 16/1024 | L | Middle Temporal Gyrus; AG | 39 | -47 | -65 | 24 | -38 | -60 | 20 | -58 | -68 | 28 |
| 16/1024 | L | Postcentral Gyrus | 7/5L | -16 | -48 | 62 | -6 | -44 | 60 | -30 | -56 | 64 |
| 15/960 | L | Lingual Gyrus | 17/18 | -6 | -58 | 2 | -2 | -48 | -1 | -10 | -64 | 8 |
| 15/960 | L | Middle Frontal Gyrus | 9 | -40 | 38 | 35 | -34 | 45 | 32 | -46 | 33 | 40 |
| 13/832 | R | Middle Temporal Gyrus | 37 | 48 | -54 | -2 | 55 | -48 | -9 | 47 | -60 | 4 |
| 12/768 | R | Cerebellum |  | 6 | -54 | -4 | 11 | -52 | -13 | 3 | -60 | 4 |
| 12/768 | R | Cingulate Gyrus | 31 | 21 | -29 | 36 | 27 | -24 | 32 | 15 | -32 | 44 |
| 12/768 | L | Cingulate Gyrus | 24 | -17 | -17 | 46 | -10 | -16 | 40 | -26 | -24 | 52 |
| 12/768 | L | Superior Frontal Gyrus | 10 | -17 | 58 | 20 | -10 | 61 | 16 | -22 | 53 | 28 |
| 11/704 | R | Cerebellum |  | 29 | -66 | -23 | 35 | -64 | -29 | 23 | -68 | -17 |
| 11/704 | R | Cerebellum |  | 7 | -61 | -37 | 15 | -60 | -41 | -6 | -64 | -37 |
| 11/704 | R | Superior Temporal Gyrus |  | 53 | -38 | 14 | 55 | -32 | 12 | 51 | -44 | 20 |
| 11/704 | R | Anterior Cingulate | 9 | 25 | 28 | 17 | 31 | 33 | 12 | 23 | 25 | 20 |
| 11/704 | R | Medial Frontal Gyrus | 10 | 1 | 57 | -10 | 3 | 61 | -17 | -6 | 53 | -5 |

**Table S2.** Final word activity locations (Figure 3 blue). See Table S1 caption for abbreviations.

|  | Region |  |  | Centre of Mass |  |  | Minimum |  |  | Maximum |  |  |
| --- | --- | --- | --- | --- | --- | --- | --- | --- | --- | --- | --- | --- |
| *Voxels/mm<sup>3</sup>* |  | Anatomical | BA | RL | AP | IS | RL | AP | IS | RL | AP | IS |
| 268/17152 | R | Superior Temporal Gyrus | 22/TE 3 | 59 | -25 | 3 | 71 | -4 | -21 | 43 | -52 | 24 |
| 192/12288 | L | Superior Temporal Gyrus | 22/TE 3 | -49 | -25 | 9 | -30 | 1 | -13 | -70 | -56 | 28 |
| 95/6080 | L | Middle Occipital Gyrus | 18 | -25 | -87 | 7 | -2 | -76 | -9 | -46 | -100 | 24 |
| 57/3648 | R | Paracentral Lobule | 4a | 4 | -41 | 63 | 19 | -24 | 48 | -18 | -52 | 80 |
| 55/3520 | R | Middle Occipital Gyrus | 18 | 26 | -87 | 13 | 39 | -80 | -5 | 11 | -96 | 24 |
| 37/2368 | R | Inferior Frontal Gyrus | 45 | 40 | 9 | 30 | 51 | 21 | 20 | 27 | 1 | 44 |
| 34/2176 | L | Middle Cingulate Cortex | 6 | -9 | -10 | 44 | -2 | 5 | 32 | -22 | -24 | 52 |
| 26/1664 | R | Postcentral Gyrus | 7/5L | 14 | -51 | 75 | 23 | -44 | 64 | 3 | -60 | 84 |
| 24/1536 | L | Middle Temporal Gyrus | 39 | -49 | -55 | 6 | -38 | -48 | -1 | -58 | -60 | 12 |
| 24/1536 | R | Thalamus |  | 18 | -27 | 16 | 27 | -20 | 8 | 11 | -32 | 28 |
| 22/1408 | R | Superior Frontal Gyrus | 6 | 7 | 22 | 53 | 11 | 33 | 44 | 3 | 13 | 64 |
| 21/1344 | R | Postcentral Gyrus | 2/3a | 29 | -30 | 36 | 39 | -24 | 28 | 19 | -36 | 44 |
| 18/1152 | L | Cerebellum |  | 2 | -35 | -9 | 11 | -28 | -13 | -6 | -44 | -1 |
| 18/1152 | L | Thalamus |  | -17 | -20 | 18 | -10 | -16 | 12 | -26 | -24 | 28 |
| 18/1152 | R | Precentral Gyrus | 6 | 57 | -2 | 47 | 63 | 9 | 36 | 55 | -12 | 52 |
| 16/1024 | L | Paracentral Lobule |  | -1 | -26 | 77 | 7 | -20 | 72 | -6 | -28 | 84 |
| 14/896 | L | Cerebellum |  | -20 | -79 | -39 | -14 | -76 | -45 | -30 | -84 | -29 |
| 13/832 | R | Middle Occipital Gyrus | 19 | 40 | -81 | 2 | 43 | -76 | -5 | 39 | -84 | 8 |
| 13/832 | R | Superior Temporal Gyrus | 41/OP 1 | 40 | -31 | 16 | 47 | -28 | 12 | 35 | -36 | 20 |
| 12/768 | L | Postcentral Gyrus | 1/5 | -32 | -42 | 71 | -26 | -36 | 68 | -38 | -48 | 76 |
| 12/768 | L | Precentral Gyrus | 6 | -29 | -23 | 64 | -22 | -20 | 60 | -34 | -28 | 68 |
| 12/768 | R | Precentral Gyrus | 6 | 39 | -6 | 29 | 43 | -4 | 24 | 31 | -12 | 32 |
| 11/704 | L | Lingual Gyrus | 18 | -8 | -80 | -9 | -6 | -76 | -13 | -10 | -88 | -1 |
| 11/704 | L | Paracentral Lobule | 4a | -10 | -42 | 70 | -6 | -36 | 64 | -14 | -48 | 76 |
| 11/704 | R | Inferior Frontal Gyrus | 45 | 44 | 27 | 23 | 51 | 29 | 20 | 39 | 25 | 28 |

**Table S3.** Activity locations supporting support vector machine (SVM) classification of high predictability filled pauses as low predictability final words (Figure 3 black outline). See Table S1 caption for abbreviations.

|  | **Region** |  |  | **Centre of Mass** |  |  | **Minimum** |  |  | **Maximum** |  |  |
| --- | --- | --- | --- | --- | --- | --- | --- | --- | --- | --- | --- | --- |
| *Voxels/mm<sup>3</sup>* |  | **Anatomical** | **BA** | **RL** | **AP** | **IS** | **RL** | **AP** | **IS** | **RL** | **AP** | **IS** |
| 125/8000 | R | Superior Temporal Gyrus | 22/TE 3 | 61 | -27 | 3 | 71 | -8 | -9 | 47 | -48 | 20 |
| 51/3264 | L | Superior Temporal Gyrus | 22/TE 3 | -57 | -23 | 7 | -46 | -8 | -1 | -66 | -36 | 20 |
| 20/1280 | L | Middle Cingulate Gyrus | 6 | -9 | -7 | 42 | -6 | 5 | 36 | -14 | -24 | 52 |
| 18/1152 | R | Calcarine Gyrus | 17 | 1 | -87 | 9 | 7 | -76 | 4 | -2 | -96 | 16 |
| 17/1088 | L | Postcentral Gyrus | 40/3b | -41 | -26 | 40 | -34 | -20 | 36 | -46 | -32 | 48 |
| 15/960 | L | Inferior Frontal Gyrus/Insula | 44/13 | -34 | 21 | 8 | -26 | 29 | 4 | -42 | 13 | 12 |
| 14/896 | R | Cuneus | 18 | 11 | -87 | 23 | 19 | -80 | 16 | 3 | -96 | 28 |
| 14/896 | R | Paracentral Lobule | 3a | 10 | -41 | 54 | 15 | -36 | 48 | 7 | -44 | 60 |
| 14/896 | R | Inferior Frontal Gyrus | 44 | 44 | 9 | 29 | 51 | 17 | 24 | 35 | 5 | 32 |

**Table S4.**
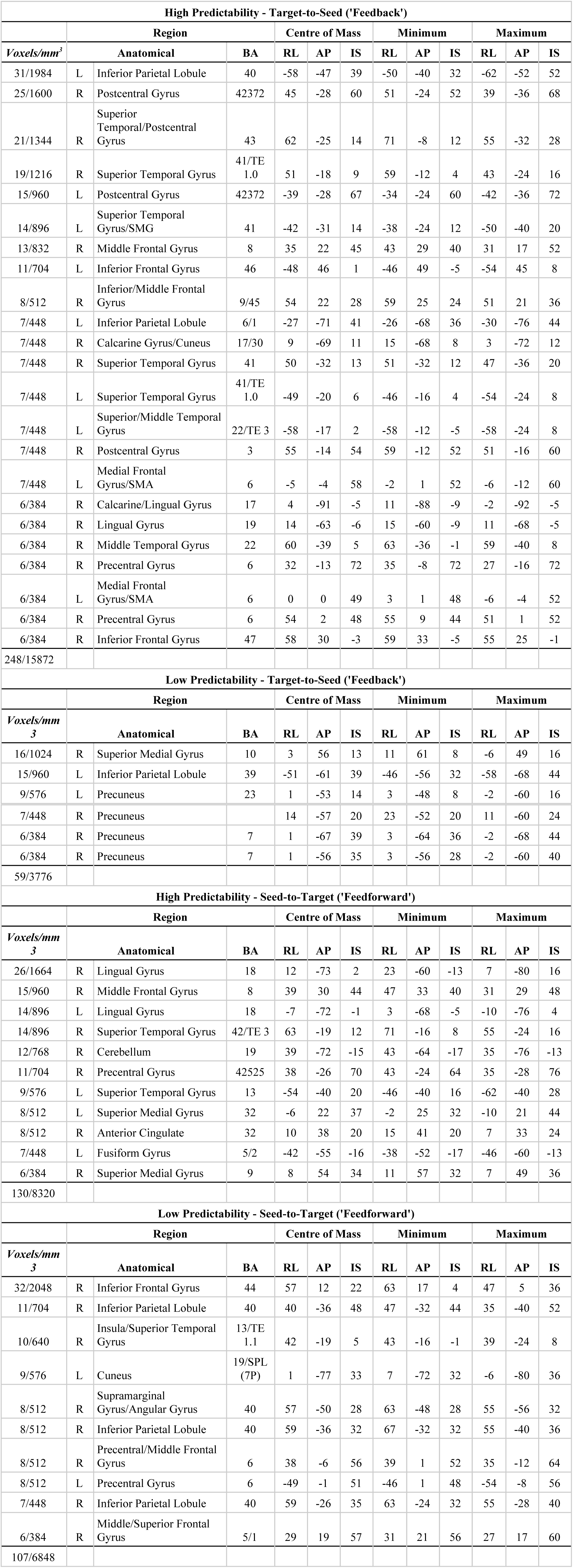
Location of network activity for greater number of seed-to-target (‘feedforward’; Figure 5 top) and target-to-seed (‘feedback’; Figure 5 bottom) connections for high- (Figure 5 red) and low-predictability (Figure 5 blue) filled pauses and final words. See Table S1 caption for abbreviations.

**Table S5.**
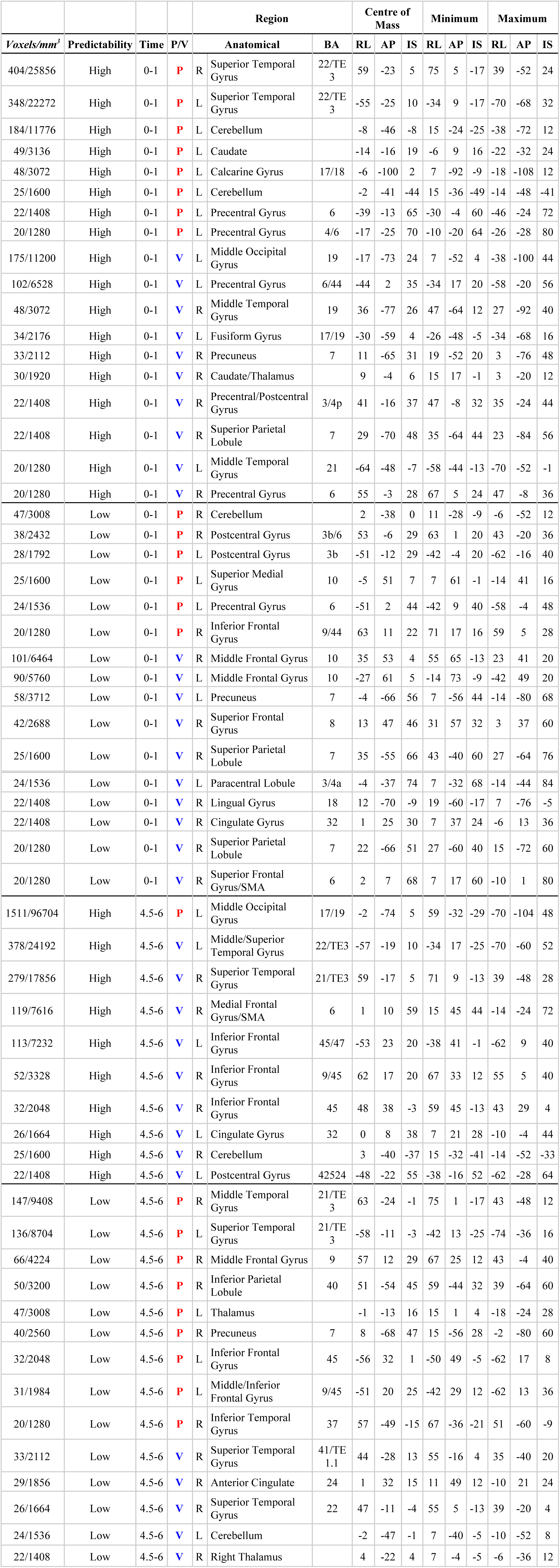
Activity locations supporting SVM classification of high and low predictability words at peaks and valleys at 0-1.5 and 4.5-6 seconds (Figure 6). Only clusters greater than or equal to 20 voxels are included. P/V = activity at peaks or valleys; see Table S1 caption for other abbreviations.

**Table S6.** Activity locations for analysis of variance (Figure S1) interaction term for factors predictability (high/low) and time (six time steps; displayed as white outline in Figure S1). Only clusters greater than or equal to 20 voxels are included. See Table S1 caption for abbreviations.

|  | Region |  |  | Centre of Mass |  |  | Minimum |  |  | Maximum |  |  |
| --- | --- | --- | --- | --- | --- | --- | --- | --- | --- | --- | --- | --- |
| *Voxels/mm<sup>3</sup>* | Anatomical |  | BA | RL | AP | IS | RL | AP | IS | RL | AP | IS |
| 441/28224 | L | Superior Temporal Gyrus | 22/TE<sub>3</sub> | -54 | -13 | 6 | -30 | 33 | -25 | -70 | -52 | 36 |
| 304/19456 | R | Superior Temporal Gyrus | 22/TE<sub>3</sub> | 59 | -16 | 1 | 71 | 9 | -21 | 35 | -40 | 24 |
| 88/5632 | R | Middle Occipital Gyrus | 19 | 37 | -75 | 12 | 55 | -60 | -5 | 19 | -92 | 36 |
| 76/4864 | L | Middle Occipital Gyrus | 19 | -42 | -74 | 8 | -34 | -60 | -5 | -54 | -92 | 20 |
| 72/4608 | R | Inferior Parietal Lobule | 40 | 45 | -51 | 54 | 59 | -40 | 36 | 27 | -72 | 64 |
| 64/4096 | R | Inferior/Middle Frontal Gyrus | 45/46 | 47 | 42 | 2 | 55 | 49 | -17 | 35 | 33 | 20 |
| 52/3328 | R | Inferior Frontal Gyrus/Precentral Gyrus | 44/9 | 49 | 4 | 30 | 59 | 29 | 24 | 43 | -12 | 40 |
| 45/2880 | L | Inferior Occipital Gyrus | 17/18 | -22 | -85 | -8 | -10 | -68 | -13 | -42 | -100 | -1 |
| 43/2752 | R | Insula | 13 | 30 | -36 | 27 | 43 | -16 | 20 | 19 | -52 | 40 |
| 28/1792 | R | Fusiform Gyrus | 19 | 31 | -61 | -13 | 43 | -44 | -21 | 23 | -72 | -9 |
| 27/1728 | R | Middle Temporal Gyrus | 21 | 65 | -49 | -3 | 75 | -36 | -13 | 55 | -60 | -1 |
| 25/1600 | R | Calcarine Gyrus | 17 | 19 | -98 | -2 | 31 | -92 | -9 | 11 | -104 | 8 |
| 24/1536 | L | Middle Occipital Gyrus | 18 | -24 | -88 | 17 | -18 | -84 | 12 | -34 | -96 | 28 |
| 23/1472 | R | Superior/Middle Temporal Gyrus | 39 | 43 | -51 | 16 | 51 | -36 | 12 | 39 | -64 | 24 |
| 22/1408 | L | Parahippocampal/Fusiform Gyrus | 19 | -26 | -58 | -8 | -22 | -48 | -17 | -34 | -68 | -1 |
| 20/1280 | L | Postcentral Gyrus | 3a | -26 | -32 | 38 | -22 | -28 | 28 | -30 | -40 | 44 |

**Figure S1.**
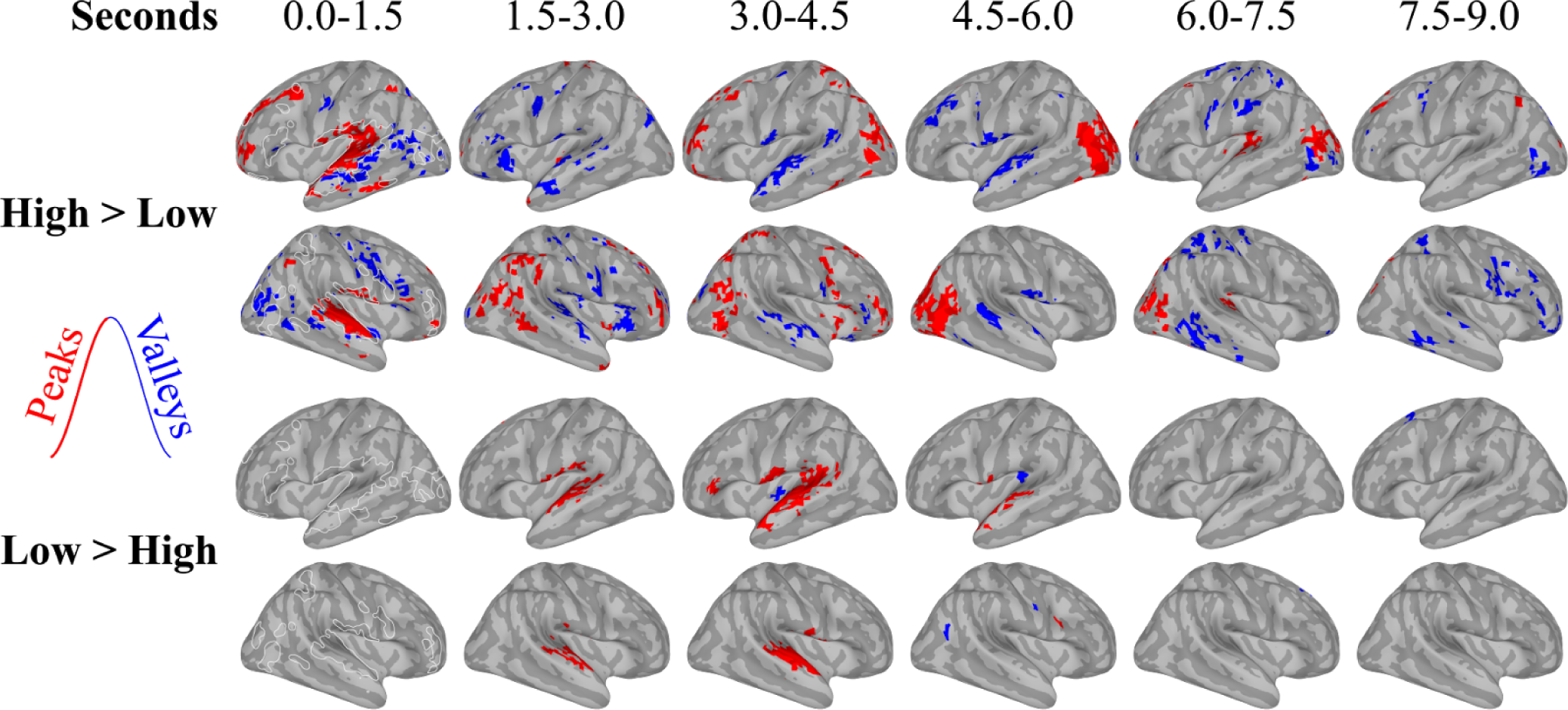
fmri study two word predictability ANOVAs. Turnpoints analysis as described in the text was done on each participant to find independent component networks corresponding to high- and low-predictability words in a television game show. A three-way repeated measures ANOVA was done on the resulting maps separately when brain responses were rising (‘Peaks’; red) and falling (‘Valleys’; blue). Displayed are second order contrasts in predictability, at fixed levels of time. The white outline on the images in the ‘0.0-1.5’ column is the interaction of time and predictability.

